# Discovery of unique mitotic mechanisms in *Paradiplonema papillatum*

**DOI:** 10.1101/2025.03.21.644664

**Authors:** Bungo Akiyoshi, Drahomíra Faktorová, Julius Lukeš

## Abstract

Diplonemids are highly diverse and abundant marine plankton with significant ecological importance. However, little is known about their biology, even in the model diplonemid *Paradiplonema papillatum* whose genome sequence is available. Examining the subcellular localization of proteins using fluorescence microscopy is a powerful approach to infer their putative function. Here we report a plasmid-based method that enables YFP-tagging of a gene at the endogenous locus. By examining the localization of proteins whose homologs are involved in chromosome segregation in other eukaryotes, we discovered several interesting features in mitotically dividing *P. papillatum* cells. Cohesin is enriched on condensed interphase chromatin. During mitosis, chromosomes organize into two rings (termed metaphase rings herein) that surround the elongating nucleolus. Homologs of chromosomal passenger complex components (INCENP, two Aurora kinases, and KIN-A), a CLK1 kinase, spindle checkpoint protein Mad1, and microtubule regulator XMAP215 localize in between the two metaphase rings, suggesting that kinetochores may assemble in between them. We also found that a homolog of the meiotic chromosome axis protein SYCP2L1 is enriched in between metaphase rings during mitosis. These features have some resemblance to the bivalent bridge (also known as the modified synaptonemal complex), which is thought to mediate the linkage between homologous chromosomes using axis components in *Bombyx mori* female meiotic metaphase I. By representing the first molecular characterization of mitotic mechanisms in *P. papillatum* and raising a number of questions, this study forms the foundation for dissecting mitotic mechanisms in diplonemids.

## Introduction

Diplonemids are highly abundant and diverse marine microorganisms (de Vargas et al., 2015; Flegontova et al., 2020; Schoenle et al., 2021) with highly flexible life strategies (Prokopchuk et al., 2022). They belong to Euglenozoa, an evolutionarily divergent group of flagellated eukaryotes that also includes kinetoplastids, euglenids, and symbiontids (Cavalier-Smith, 2016; Kostygov et al., 2021). Dissecting the biology of diplonemids is key for understanding their ecological importance as well as the origin of distinct biological processes and pathogenicity in Euglenozoa, such as trypanosomes, leishmanias and other serious pathogens. As exemplified by their membrane-trafficking machinery, the gene-rich diplonemids possess both unique and conserved proteins, proteins previously considered as kinetoplastid-specific, as well as those with sporadic distribution across eukaryotes (Záhonová et al., 2025).

So far, very little is known about the proliferation mechanisms in diplonemids. Classic electron microscopy studies showed that diplonemids have chromosomes that are condensed even in interphase and undergo closed mitosis with intact nucleolus (Porter, 1973; Triemer and Ott, 1990; Triemer, 1992). As cells enter mitosis, the nucleolus gets elongated by an unknown mechanism. It has been reported that chromosomes form one distinct ring surrounding the elongating nucleolus, which then separate as two rings in anaphase (Porter, 1973; Triemer, 1992). Interestingly, a higher number of spindle microtubules was observed between the separating chromosomal rings rather than between the rings and the poles (Triemer, 1992). Although many mitotic proteins involved in chromosome organizations in other eukaryotes are conserved in diplonemids, they have not been characterized yet. The nuclear genome of the model diplonemid *Paradiplonema papillatum* is ∼280 Mb with ∼32,000 protein-coding genes (Valach et al., 2023). The size of chromosomes is 1.1–1.8 Mb and it is estimated that there are ∼180 chromosomes. It remains unknown how these numerous chromosomes organize into a ring-like structure during mitosis.

Kinetochores are the macromolecular complex that drives chromosome segregation during mitosis and meiosis in eukaryotes (Musacchio and Desai, 2017). Although components of kinetochores are widely conserved among eukaryotes (Drinnenberg and Akiyoshi, 2017; van Hooff et al., 2017), interesting exceptions are found in Euglenozoa. While euglenids have canonical kinetochore proteins (Ebenezer et al., 2019; Butenko et al., 2020), kinetoplastids have a unique set of kinetochore proteins that are so far exclusively found in this group (Akiyoshi and Gull, 2014). It remains unclear when and how the unique kinetoplastid kinetochore system evolved (Akiyoshi, 2016). Because phylogenetic analysis places kinetoplastids next to diplonemids rather than euglenids (Butenko et al., 2020; Lax et al., 2021), it is important to understand diplonemid kinetochores to gain hints to this question. Interestingly, it remains unclear what kind of kinetochore proteins are present in diplonemids (Butenko et al., 2020; Benz et al., 2024). Although bioinformatics search identified homologs of some kinetoplastid kinetochore proteins, such as CLK kinases (involved in splicing) and SYCP2 and SYCP3 (involved in meiotic synapsis), these broadly conserved proteins are known to have non-kinetochore functions outside of kinetoplastids (Butenko et al., 2020; Benz et al., 2024). Using *P. papillatum* for which genome sequence and transfection methods are available (Faktorová et al., 2020; Valach et al., 2023), we previously found that a putative CENP-A homolog showed dot signals in interphase cells (Benz et al., 2024). However, without any kinetochore marker available, we could not confirm that the dots represent kinetochore signals. Furthermore, due to difficulties in performing immunofluorescence microscopy in this organism, such as its poor stickiness to glass coverslips, it was not possible to determine the localization of the putative CENP-A or other proteins in mitotic cells. To overcome these issues, here we developed an endogenous YFP-tagging method and determined mitotic localization for proteins whose homologs play mitotic/meiotic roles in other eukaryotes.

## Results

### Cell cycle analysis by DAPI staining

To gain insights into the mechanism of chromosome segregation in *P. papillatum*, we first performed DAPI staining analysis on logarithmically growing cells. Due to poor stickiness to microscope glass slides or coverslips, we could not immobilize *P. papillatum* cells using a standard protocol. We therefore fixed cells in solution, resuspended them in a small volume of mounting media with DAPI, and mounted cells onto a slide. This simple modification allowed us to image many cells by fluorescence microscopy (Figure 1a). By imaging >1000 cells in three independently growing cultures (Figure 1b–d, Table S1), we found that the vast majority of cells was non-mitotic (97.6 ± 0.5%). As observed by electron microscopy studies (Porter, 1973; Triemer and Ott, 1990; Tashyreva et al., 2018, 2023, 2025), chromosomes are condensed even in interphase and are visible as sausage-like structures (Figure 1b). We also observed cells of unknown cell cycle stage with crescent-shape DNA morphology (0.65 ± 0.11%) as well as few anucleate cells (0.06 ± 0.11%). In a minority of the population (1.6 ± 0.45%), we observed cells undergoing mitosis. The following categories of mitotic stages were observed based on their DNA morphology: 1) metaphase with one metaphase ring (0.06 ± 0.06%), 2) “putative metaphase” with two metaphase rings whose DNA intensity peaks were separated by up to 1 µm (0.45 ± 0.29%), and 3) late anaphase when DNA is separated further and do not appear as rings (1.1 ± 0.24%). In this experiment, we did not observe cells that were in early anaphase (two rings that are separated by >1 µm), presumably because they are even rarer. In the category 1 and 2, the observed rings surrounded the elongating nucleolus (Figure 1c), as previously reported (Porter, 1973; Triemer, 1992). Based on the fact that we observed cells with two rings much more frequently than those with one ring, we speculate that the cells with two rings are likely in metaphase, a possibility supported by the localization of two mitotic proteins whose signals are lost in anaphase (see below). However, if this is the case, these observations raise a number of questions: How do numerous chromosomes organize into two rings? How are duplicated chromosomes linked together (assuming that the two metaphase rings correspond to two sets of duplicated sister chromatids)? Where do kinetochores assemble on the metaphase rings? How do spindle microtubules interact with the rings? How do they achieve bi-oriented attachments?

**Figure 1.**
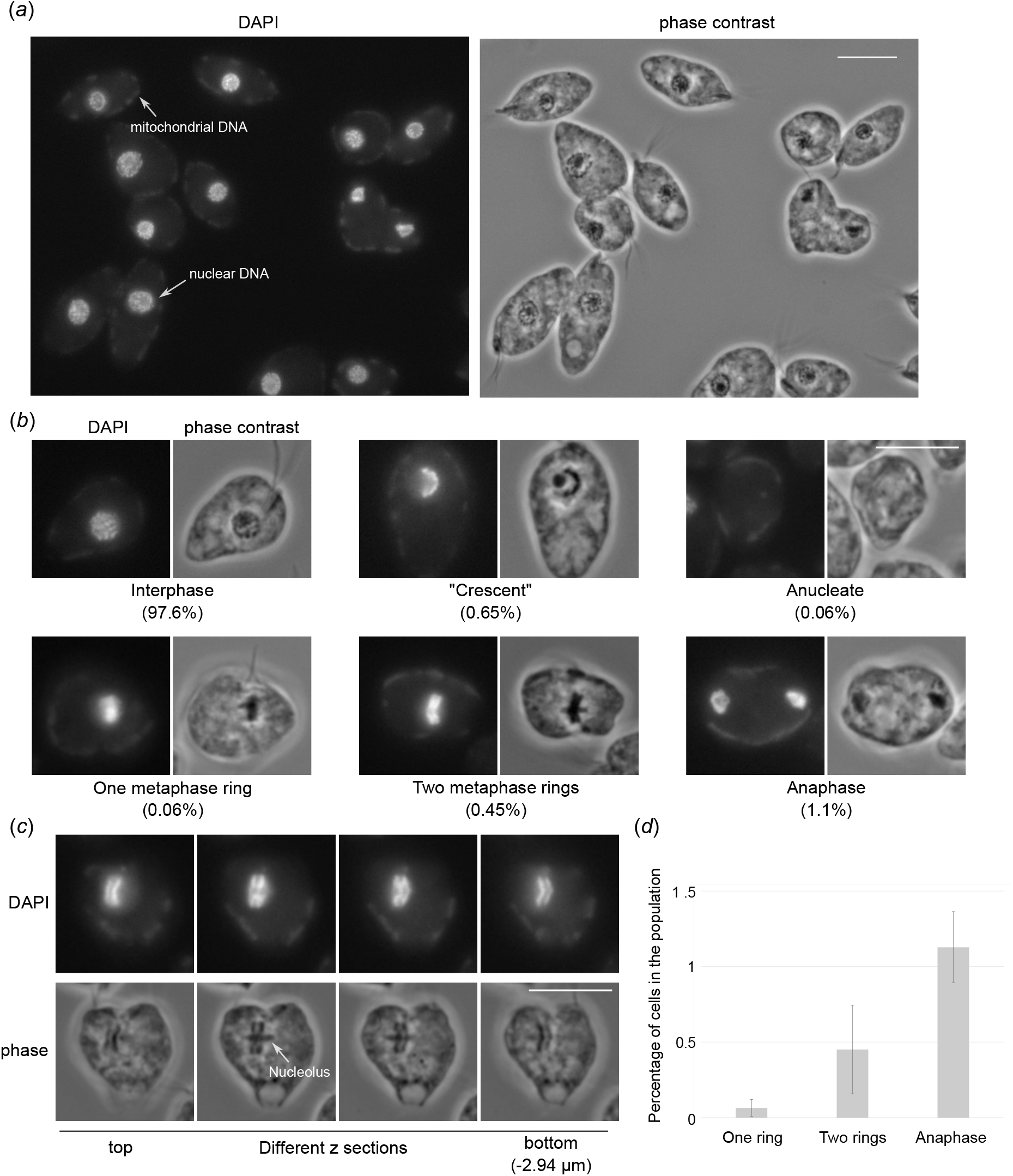
Quantification of cell cycle analysis by DAPI staining. (a) A wide field of view of *Paradiplonema papillatum* cells stained with DAPI. Bars, 10 µm. (b) Example of cells with distinct DNA morphology and their quantification. More than 1000 cells were analyzed (N=3). For raw data, see Table S1. (c) An example of a cell that has two metaphase rings. Different z sections are shown. (d) Quantification of mitotic cells in the population. Error bars stand for standard deviation.

### C-terminal YFP-tagging method enables examination of mitotic cells in *P. papillatum*

To gain insights into the above questions, we aimed to systematically examine the localization of various conserved mitotic proteins by fluorescence microscopy. In *P. papillatum*, chromosomal integration of a DNA fragment can be achieved via homologous recombination by using ∼1.5 kb homology arms (Kaur et al., 2018; Faktorová et al., 2020). Using this method, *P. papillatum* genes have been tagged with protein A or V5 (Faktorová et al., 2020, 2023). We adapted the system to enable YFP tagging using a plasmid-based method, similar to the one established for *Trypanosoma brucei* (Kelly et al., 2007). For C-terminal YFP tagging, two ∼2 kb homology arms are amplified from genomic DNA (Figure 2a, Table S2). The first corresponds to a 2 kb DNA fragment downstream of the open reading frame, starting just after the stop codon. The second is a fragment that starts from 2 kb upstream of the stop codon and ends just prior to the stop codon so that the open reading frame is in frame with YFP. These two DNA fragments are cloned into a vector (pBA3294 or pBA3235: YFP with a neomycin resistant marker). A unique restriction enzyme site is introduced in between the two DNA fragments (typically *Not*I), which is used to linearize the plasmid to enable C-terminal YFP tagging of a gene at the endogenous locus. pBA3295 allows tdTomato tagging with a hygromycin selection marker. After electroporation and drug selection, a population of transgenic cells are obtained after ∼10 days.

**Figure 2.**
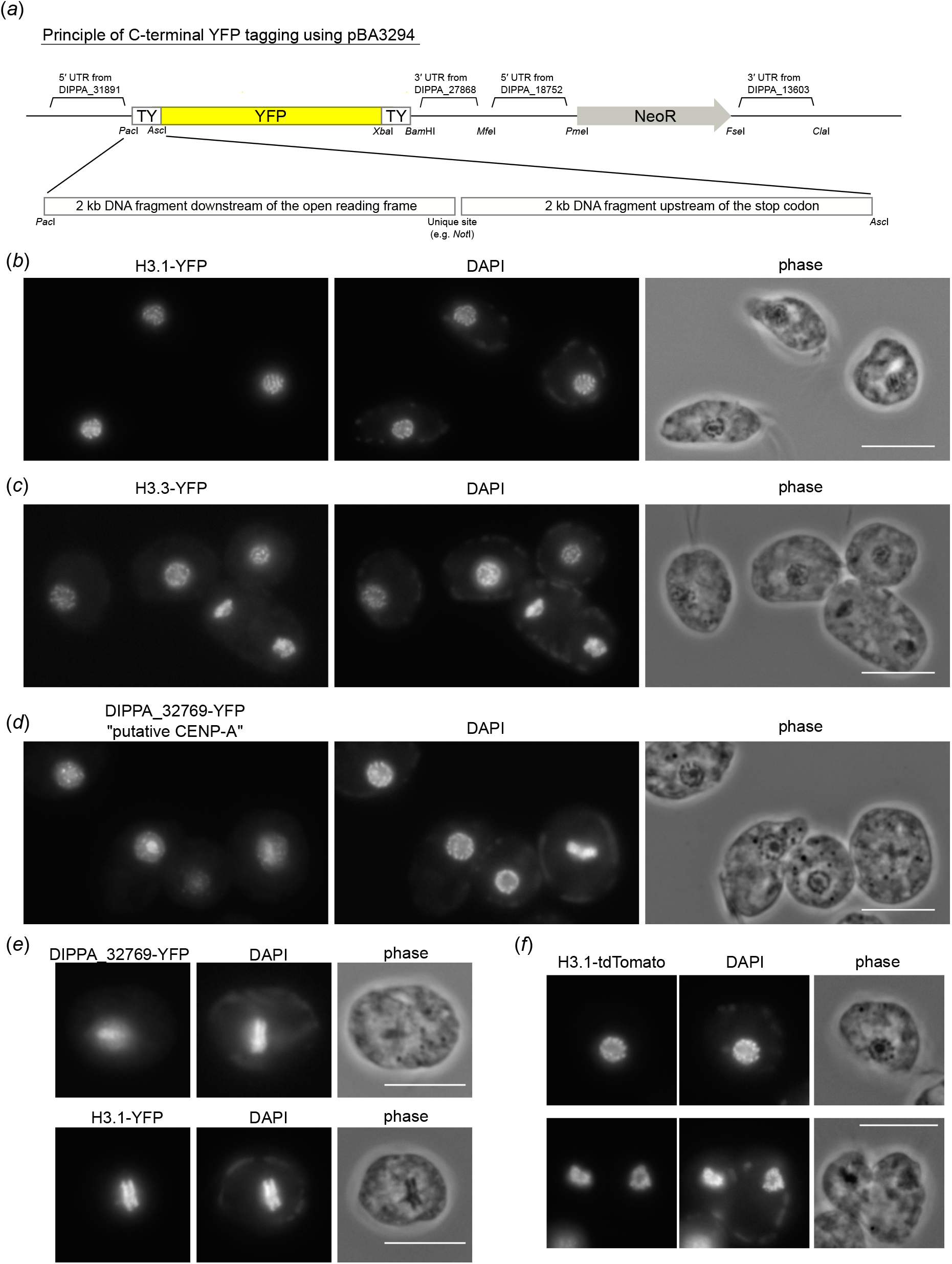
The putative CENP-A homolog does not show kinetochore localization. (a) Schematic of the YFP-tagging vector, pBA3294. For C-terminal tagging of a gene of interest, two homology arms are inserted into *Pac*I and *Asc*I sites of pBA3294. pBA3235 uses DIPPA_28792’s 5′ UTR for YFP as well as different restriction sites. pBA3295 has tdTomato and hygromycin resistant gene instead of YFP and neomycin resistant gene of pBA3294. Full sequences of these vectors are shown in Table S2. (b) H3.1-YFP shows chromatin signal. Bars, 10 µm. (c) H3.3-YFP shows chromatin signal. (d) The putative CENP-A homolog-YFP shows dots in the nucleus. (e) Putative CENP-A-YFP does not localize on chromatin in metaphase, while H3.1-YFP does. (f) H3.1-tdTomato shows chromatin signal.

Using this method, we examined the localization of a histone H3-like protein H3.1 (DIPPA_26288) and found chromatin signal for both YFP- and tdTomato-fusion proteins (Figure 2b, 2f). Another histone H3-like protein H3.3 (DIPPA_21362) also showed chromatin signal (Figure 2c). The putative centromeric H3 variant (DIPPA_32769) has sequence features characteristic for CENP-A and forms dots in interphase cells (Benz et al., 2024). Although dot signals were found for DIPPA_32769-YFP in interphase cells (Figure 2d) as reported previously, the signals were not on metaphase rings during mitosis (Figure 2e). By contrast, H3.1-YFP signals were on metaphase rings. These results do not support the possibility that DIPPA_32769 is a kinetochore protein. However, we cannot exclude a possibility that fusing YFP interfered with its proper localization.

### Kinetoplastid-like CPC compositions in *P. papillatum*

To gain insights into mitotic mechanisms in *P. papillatum*, we next examined the localization of INCENP (DIPPA_09943), a component of the chromosomal passenger complex (CPC) that localizes near kinetochores in metaphase and then at central spindles in anaphase in many eukaryotes including kinetoplastids (Cooke et al., 1987; Li et al., 2008). DAPI signals were used to screen rare mitotic cells. We found that *P. papillatum* INCENP localizes in between metaphase rings and then at central spindles in anaphase (Figure 3a), thereby showing that it is a genuine CPC subunit. We next examined the localization of four Aurora homologs to identify which Aurora kinase(s) are the catalytic subunit of the *P. papillatum* CPC. Interestingly, Aurora1 (DIPPA_00804) and Aurora2 (DIPPA_18318) showed an INCENP-like localization pattern (Figure 3b and 3c). In contrast, Aurora3 (DIPPA_20993) had a nuclear signal (Figure 3d), and Aurora4 (DIPPA_24328) localized near basal bodies (Figure 3e). These localization patterns are somewhat similar to *T. brucei* AUK2 that has nuclear signal (Stortz et al., 2017) and AUK3 that localizes near basal bodies (Akiyoshi, 2020). In addition, we found that a kinesin-like protein (DIPPA_28866) that has the highest similarity to KIN-A (a CPC subunit in *T. brucei* (Ballmer and Akiyoshi, 2024)) had a CPC-like localization pattern (Figure 3f), raising a possibility that the CPC in *P. papillatum* is compositionally similar to that in kinetoplastids.

**Figure 3.**
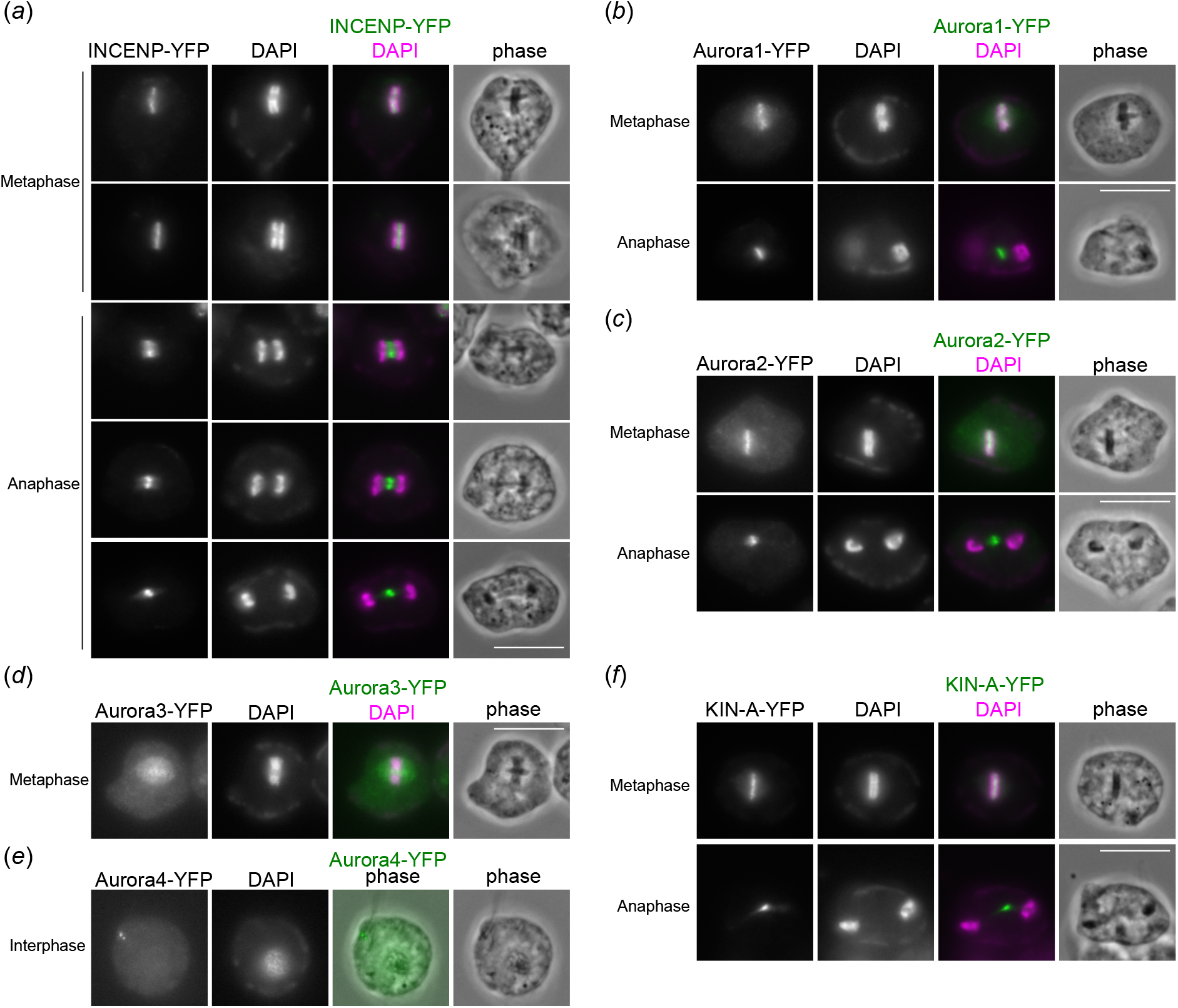
CPC in *P. papillatum* contains INCENP, two Aurora kinases, and a KIN-A homolog. (a) INCENP-YFP shows dynamic localization pattern during mitosis. Bars, 10 µm. (b) Aurora1-YFP shows a typical CPC localization pattern. (c) Aurora2-YFP also shows CPC localization. (d) Aurora3-YFP shows a diffuse nuclear signal during metaphase. (e) Aurora4-YFP localizes at basal bodies. (f) KIN-A-YFP shows a typical CPC localization pattern.

### Homologs of regulatory kinetoplastid kinetochore proteins localize in between metaphase rings

We next examined the localization of broadly conserved proteins that are thus far known to have kinetochore functions only in kinetoplastids. CLK kinases regulate kinetochore functions in kinetoplastids (called KKT10 and KKT19 in *T. brucei*) (Ishii and Akiyoshi, 2020; Saldivia et al., 2020), while they are involved in RNA splicing functions in other eukaryotes (Corkery et al., 2015). We found that CLK1 (DIPPA_05595) localized in between the metaphase rings and some cytoskeletal structures in *P. papillatum* (Figure 4a). Interestingly, the signal disappeared from the rings in anaphase, reminiscent of the localization pattern of KKT10 and KKT19 kinetochore proteins in *T. brucei* (Akiyoshi and Gull, 2014). It is therefore possible that this signal “in between metaphase rings” represents kinetochore localization in *P. papillatum*. Given that KKT10 and KKT19 similarly disappear from kinetochores in anaphase in *T. brucei*, another corollary is that the cells that have two metaphase rings are in metaphase even though the two rings have significant distance in between.

**Figure 4.**
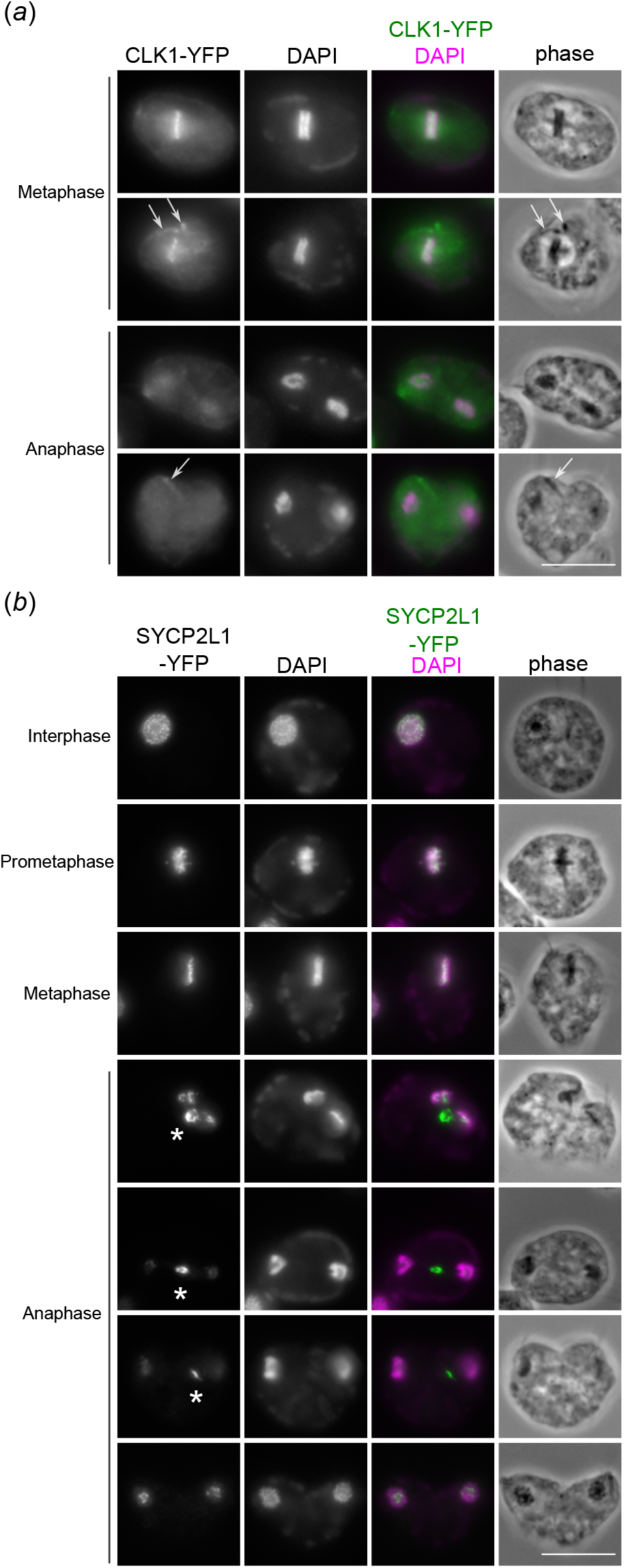
CLK1 and SYCP2 localize in between metaphase rings. (a) CLK1-YFP localizes in between two metaphase rings, but disappears from chromosomes in anaphase. It also has cytoskeleton signals (arrows), derived from flagella, basal bodies and/or papilla. Bars, 10 µm. (b) SYCP2L1-YFP localizes on interphase chromatin. During mitosis, the signals organize into chromosomal rings during putative prometaphase and localizes in between the metaphase rings. During anaphase, the signal is found in the inner side (i.e. towards the center) of each ring as well as in between separated nuclei (asterisks).

SYCP2 homologs localize at mitotic kinetochores in kinetoplastids (called KKT17 and KKT18 in *T. brucei*), while they are used as components of the chromosome axis and synaptonemal complex during meiosis in many eukaryotes (possibly including kinetoplastids) (Tromer et al., 2021; Adams and Davies, 2023). We found that a SYCP2-like protein called SYCP2L1 (DIPPA_35871), which localized on chromatin in interphase, occurred in between the metaphase rings (Figure 4b), suggesting that, like KKT17 and KKT18 in *T. brucei*, SYCP2L1 may have kinetochore functions in *P. papillatum*. Besides chromatin-proximal signals, SYCP2L1 also showed an interesting signal of an unknown location in anaphase cells, which differs from CPC’s central spindle signal (Figure 4b).

### A putative Mad1 homolog and XMAP215 localize in between metaphase rings

We next made YFP-fusions for putative homologs of widely conserved proteins that localize at kinetochores in many organisms, but not in kinetoplastids. Spindle checkpoint proteins localize at kinetochores to monitor kinetochore-microtubule attachments (Musacchio and Salmon, 2007). We found that a spindle checkpoint Mad2 homolog (DIPPA_17925) localizes near basal bodies and feeding apparatus (Figure 5a), the former localization pattern being reminiscent of Mad2 in *T. brucei* (Akiyoshi and Gull, 2013; Billington et al., 2023). By contrast, a putative homolog of another spindle checkpoint protein Mad1 (DIPPA_30775) localized at nuclear pores during interphase and in between metaphase rings during metaphase, while the signal largely disappeared from chromosomes in anaphase (Figure 5b). Mad1 also showed signals at putative spindle pole areas. A microtubule regulator XMAP215 (DIPPA_00331) localizes at kinetochores in metaphase and anaphase in some organisms (Miller et al., 2016; Herman et al., 2020) (Figure 5c). We found that it also localized in between metaphase rings in metaphase, while central spindle and spindle midzone-like signals were observed in early anaphase.

**Figure 5.**
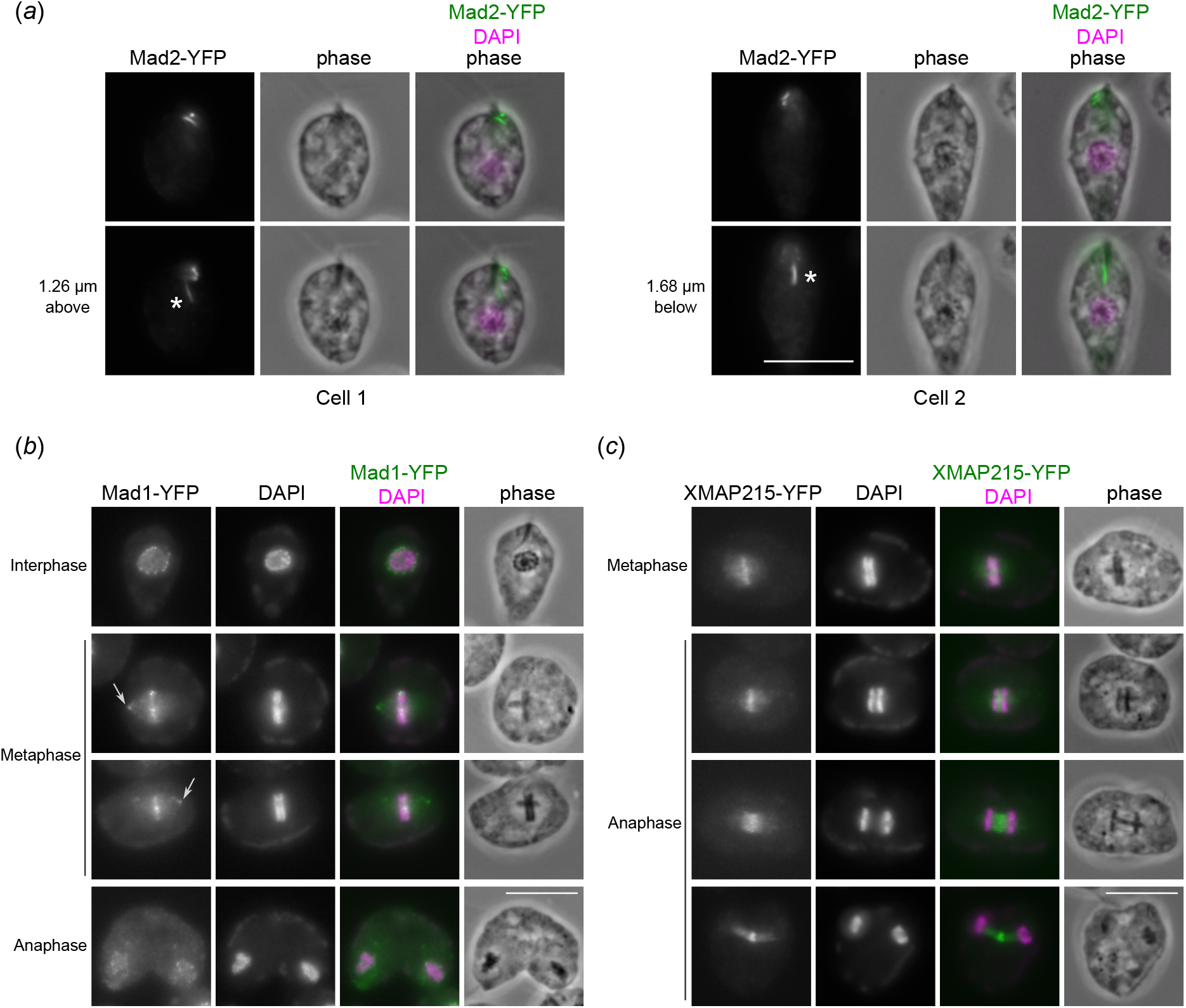
Mad1 and XMAP215 localize in between metaphase rings. (a) Mad2-YFP localizes near basal bodies and feeding groove (asterisks). Bars, 10 µm. (b) Mad1-YFP shows nuclear pore signals in interphase. During metaphase, the signal is found in between the metaphase rings as well as putative spindle pole areas (arrows). (c) XMAP215-YFP localizes in between metaphase rings, then shows signal at central spindles and spindle midzone.

### Cohesin localizes on interphase and mitotic chromatin, not in between metaphase rings

In eukaryotes, duplicated sister chromatids are linked together by the cohesin complex (Yatskevich et al., 2019), while the condensin complex plays major roles in chromosome organization (Hirano, 2012). To examine the mechanism of chromosome organization in *P. papillatum*, we examined cohesin subunits SMC1 (DIPPA_30927) and SCC3 (DIPPA_15505), as well as a condensin subunit SMC4 (DIPPA_23430). We found that cohesin subunits were enriched on interphase chromatin (Figure 6a and 6b). By contrast, condensin SMC4 showed mostly diffuse nuclear signal in interphase cells (Figure 6c). Interestingly, both cohesin and condensin were enriched on chromatin in metaphase with no obvious enrichment in between the metaphase rings (Figure 6a-c).

**Figure 6.**
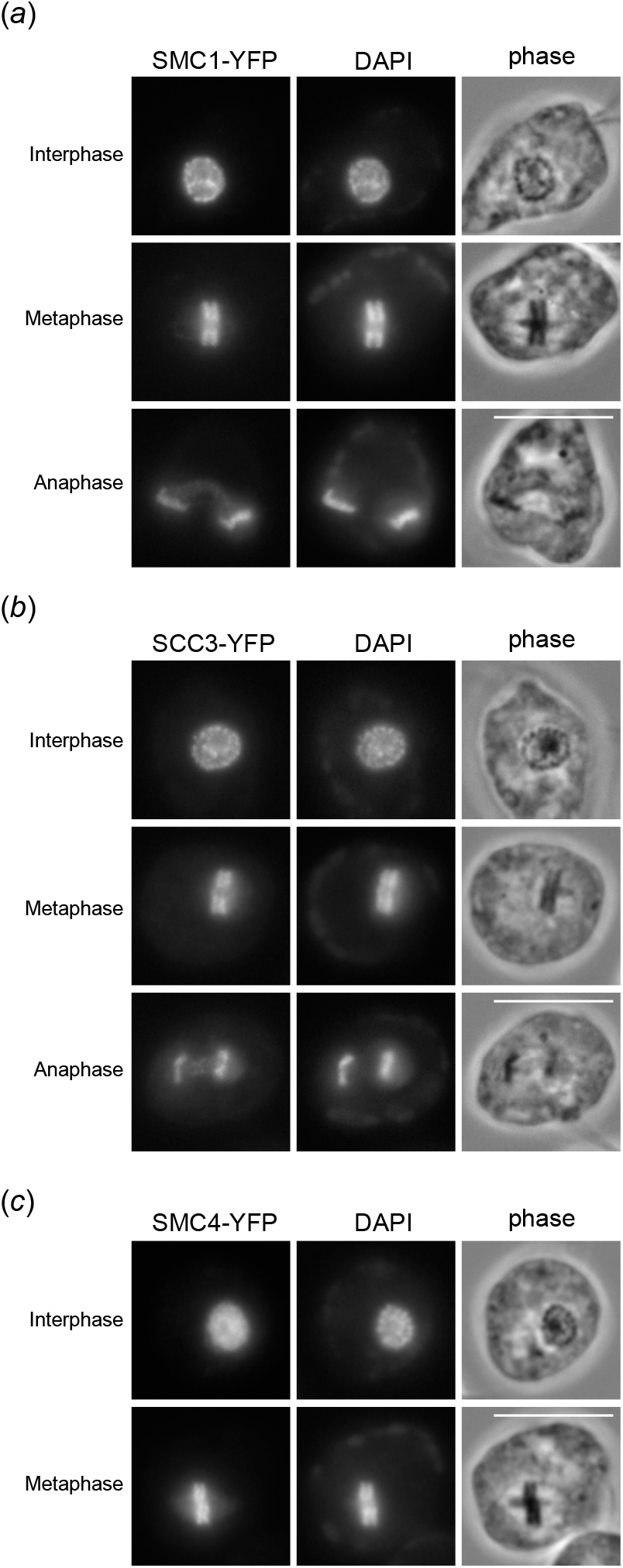
Cohesin subunits are enriched on chromatin, but not in between the metaphase rings. (a) A cohesin subunit SMC1-YFP shows chromatin signal in interphase, metaphase, and anaphase. No obvious enrichment was observed in between metaphase rings. During anaphase, a diffuse nuclear signal was observed for SMC1 and SCC3. Bars, 10 µm. (b) SCC3-YFP shows similar localization pattern as SMC1. (c) A condensin subunit SMC4-YFP shows a diffuse nuclear signal in interphase and chromatin signal in metaphase.

## Discussion

Through YFP-tagging of genes, we have revealed several interesting features of chromosome biology in *P. papillatum*. Our finding that condensed interphase chromatin has cohesin, not condensin, suggests that cohesin may be responsible for the interphase chromosome organization in diplonemids. In mammalian cells, depletion of Wapl causes stable chromosomal association of cohesin, leading to chromatin condensation in interphase cells and chromosome mis-segregation in mitosis (Tedeschi et al., 2013). It will be interesting to understand how diplonemids cope with permanently condensed chromosomes.

One surprising finding from this study is the two metaphase rings with significant space in between. Presence of CLK1 and Mad1 signals on these chromosomes supports the possibility that these cells are prior to anaphase but raises a number of questions including how chromosomes are linked. Conservation of cohesin subunits and regulators including the Eco1 acetyltransferase (DIPPA_13679) and putative SMC3 acetylation sites (K109 and K110 in DIPPA_26840) supports the idea that sister chromatid cohesion is mediated by cohesin complexes in diplonemids, which get cleaved by separase (DIPPA_16480) at the onset of anaphase. Although we did not observe strong cohesin signals in between the metaphase rings, it is possible that a low amount of cohesin is sufficient to hold sister chromatids together. An alternative possibility is that something else holds sister chromatids together during mitosis. In female meiosis of *Bombyx mori* that lacks chiasmata, homologous chromosomes are separated by ∼700 nm but are connected by a structure called the bivalent bridge that includes SYCP2, HOP1, and PCH2 (Xiang et al., 2024). It will be interesting to examine if HOP1 and PCH2 localize near metaphase rings in mitotic *P. papillatum* cells. Furthermore, we formally cannot exclude the possibility that the two metaphase rings do not correspond to duplicated sister chromatids. For example, it might be possible that one set of sister chromatids (both located within one ring) may pair up with another set of sisters (within the other ring), which could be a homologous chromosome (although *P. papillatum* is thought to be haploid). However, this would mean that sister chromatids do not separate from each other during mitosis, a phenomenon never reported in eukaryotes. Further investigation is necessary to reveal the nature of metaphase rings.

The CPC localizes at metaphase kinetochores in almost all studied eukaryotes, including kinetoplastids (Hochegger et al., 2013). The most conserved components of the CPC are the Aurora B kinase and its activator INCENP (van Hooff et al., 2017). While recruitment of the CPC to kinetochores relies on Survivin and Borealin in many eukaryotes (Komaki et al., 2022), in kinetoplastids it relies on a unique component called KIN-A (Ballmer and Akiyoshi, 2024). Our finding that a KIN-A homolog is a CPC component in *P. papillatum* is consistent with the close evolutionary relationship between diplonemids and kinetoplastids (together called glycomonads) (Cavalier-Smith, 2016), which is further supported by the localization of CLK1 and SYCP2L1 in mitotic chromosomes. In this sense, chromosome segregation machinery in *P. papillatum* appears to resemble that in kinetoplastids. However, we also found that a Mad1 homolog localizes in between metaphase rings. The function of Mad1 remains unclear because its canonical interaction partner Mad2 localizes at a different location in *P. papillatum*. In *T. brucei*, Mad2 and its interaction partner MBP65 localize at microtubule quartet near basal bodies (Akiyoshi, 2020; Billington et al., 2023). Mad2 shows similar localization pattern in *P. papillatum*, and MBP65 homologs are present in diplonemids (DIPPA_21359 in *P. papillatum*). It will be important to dissect the function of Mad1, Mad2, and MBP65, which could provide insights into the evolutionary origin of the spindle checkpoint mechanism in eukaryotes.

Mechanisms of chromosome bi-orientation and segregation in *P. papillatum* remains unknown. The CPC and XMAP215, which play key roles in error correction in other eukaryotes (Lampson and Cheeseman, 2011; Miller et al., 2016), localize in between metaphase rings, implying their possible roles in promoting bi-orientation in diplonemids. During anaphase, both the CPC and XMAP215 localize at central spindles. Together with previous observations by electron microscopy that there are a lot of microtubules in between the two separating rings in anaphase (Triemer, 1992), these data could imply that chromosomes are pushed apart in anaphase, a phenomenon observed in some eukaryotes (Laband et al., 2017; Yu et al., 2019; Chen et al., 2024). It will be important to understand what drives chromosome segregation in diplonemids.

In conclusion, our study starts to provide molecular insights into mechanisms of chromosome organization and segregation in *P. papillatum*. It is important to note that there is still no structural kinetochore protein identified in diplonemids, leaving the intriguing possibility open that diplonemids

have a hitherto unknown type of kinetochores. In *T. brucei*, CLK1 and SYCP2 homologs co-purify with structural kinetochore components (Akiyoshi and Gull, 2014), while Mad1 co-purifies with kinetochore proteins in yeast (Akiyoshi et al., 2010). It is therefore possible that immunoprecipitation coupled with mass spectrometry for these homologs could identify structural kinetochore proteins in *P. papillatum*.

## Materials and methods

### Tagging vectors, plasmids, and primers

Sequences of primers and tagging vectors are provided in electronic supplementary material Table S2. pBA3235 (first generation YFP-tagging vector), pBA3294 (second generation YFP-tagging vector) and pBA3295 (tdTomato-tagging vector) were synthesized by GeneArt (Thermo Fisher). For C-terminal tagging using pBA3294 (YFP) or pBA3295 (tdTomato), *Pac*I and *Asc*I restriction sites were used to insert two ∼2 kb homology arms that were amplified from genomic DNA by PCR using KOD one polymerase (Merck). Primers were designed using the NEBuilder assembly tool (New England Biolabs), avoiding repetitive sequences if necessary and possible. One unique site was introduced in between the two fragments (typically *Not*I; if *Not*I was not unique, different enzymes were used): the first fragment corresponding to downstream of the open reading frame of the gene (starting just after its stop codon) surrounded with *Pac*I and *Not*I restriction sites, and the second fragment corresponding to the 2 kb DNA fragment starting from 2kb upstream of the stop codon and ending just before the stop codon) surrounded with *Not*I and *Asc*I. After cutting the fragments with respective restriction enzymes, the two DNA fragments were ligated into pBA3294 or pBA3295 that were cut with *Pac*I and *Asc*I. For C-terminal tagging using pBA3235, *Sbf*I and *Not*I restriction sites were used in combination with *Fse*I. Plasmids were screened and validated by nanopore whole plasmid sequencing (Plasmidsaurus). We occasionally detected mismatches between nanopore sequencing results and expected plasmid sequences which were made *in silico* based on the genome sequence, especially in repetitive sequences in 3′UTR regions. However, we did not attempt to figure out whether they are errors of nanopore sequencing or genome sequence.

### Diplonema culture and transfection

All cell lines used in this study were derived from *Paradiplonema papillatum* (ATCC 50162). Cells were grown at 27 °C in liquid medium containing 36 g/L Instant Ocean Sea Salt (Instant Ocean), 1 g/L trypton (Formedium, TRP01), and 1% fetal bovine serum (Merck, F9665) in vented flasks.

Five to 10 µg plasmids were linearized by *Not*I or other enzymes, followed by ethanol precipitation. DNA was resuspended in 20 µL of transfection reagent (Ingenio Electroporation Kit for the EZporator Electroporation System, Cambridge Biosciences) and transfected into ∼3 x 10^7^ cells using Amaxa Nucleofector IIb (Lonza Bioscience). Transfected cells were selected by the addition of 75 µg/mL G418 (pBA3235 and pBA3294 derivatives) or 125 µg/mL hygromycin (pBA3295 derivatives) (Merck). All cell lines used in this study are listed in electronic supplementary material Table S2.

### Microscopy

Cells were pelleted by centrifugation at 1300 g for 5 min and fixed by 4% formaldehyde solution diluted in PBS (Life technologies, #28906) for 5 min. Cells were washed by 1 mL PBS twice, resuspended in a small volume of DABCO mounting media (1% w/v 1,4-diazabicyclo[2.2.2]octane, 90% glycerol, 50 mM sodium phosphate pH 8.0) with 100 ng/mL DAPI, and mounted onto glass slides. Images were captured on an Axioimager.Z2 microscope (Zeiss) installed with ZEN using a Hamamatsu ORCA-Flash4.0 camera with 63x objective lenses (1.40 NA). Typically, 15–30 z sections spaced 0.24 μm apart were collected. Images were analyzed in ImageJ/Fiji (Schneider et al., 2012). Figures were made in Inkscape (version 1.3, https://inkscape.org/).

## Acknowledgments

We acknowledge support from a Wellcome Discovery Award (227243/Z/23/Z to B.A.) and the Czech Grant Agency 23-06479X (to J.L.). We also thank Sam Taylor and Dipika Mishra for comments on the manuscript.

## Author contribution

B.A. carried out all the experiments, analyzed the data, and wrote the manuscript. D.F. and J.L. provided *Paradiplonema papillatum* cells and advice on genetic manipulation, and edited the manuscript. All authors gave final approval for publication and agreed to be held accountable for the work performed therein.

## Rights retention

This research was funded in whole, or in part, by Wellcome Trust [227243/Z/23/Z]. For the purpose of open access, the author has applied a CC BY public copyright licence to any Author Accepted Manuscript version arising from this submission.

## References

Adams, I.R., and O.R. Davies. 2023. Meiotic chromosome structure, the synaptonemal complex, and infertility. Annu Rev Genomics Hum Genet. 24:35–61. doi:10.1146/annurev-genom-110122-090239.

Akiyoshi, B. 2016. The unconventional kinetoplastid kinetochore: from discovery toward functional understanding. Biochem. Soc. Trans. 44:1201–1217. doi:10.1042/BST20160112.

Akiyoshi, B. 2020. Analysis of a Mad2 homolog in Trypanosoma brucei provides possible hints on the origin of the spindle checkpoint. 2020.12.29.424754. doi:10.1101/2020.12.29.424754.

Akiyoshi, B., and K. Gull. 2013. Evolutionary cell biology of chromosome segregation: insights from trypanosomes. Open Biol. 3:130023. doi:10.1098/rsob.130023.

Akiyoshi, B., and K. Gull. 2014. Discovery of unconventional kinetochores in kinetoplastids. Cell. 156:1247–1258. doi:10.1016/j.cell.2014.01.049.

Akiyoshi, B., K.K. Sarangapani, A.F. Powers, C.R. Nelson, S.L. Reichow, H. Arellano-Santoyo, T. Gonen, J.A. Ranish, C.L. Asbury, and S. Biggins. 2010. Tension directly stabilizes reconstituted kinetochore-microtubule attachments. Nature. 468:576–579. doi:10.1038/nature09594.

Ballmer, D., and B. Akiyoshi. 2024. Dynamic localization of the chromosomal passenger complex in trypanosomes is controlled by the orphan kinesins KIN-A and KIN-B. Elife. 13:RP93522. doi:10.7554/eLife.93522.

Benz, C., M.W.D. Raas, P. Tripathi, D. Faktorová, E.C. Tromer, B. Akiyoshi, and J. Lukeš. 2024. On the possibility of yet a third kinetochore system in the protist phylum Euglenozoa. mBio. 15:e0293624. doi:10.1128/mbio.02936-24.

Billington, K., C. Halliday, R. Madden, P. Dyer, A.R. Barker, F.F. Moreira-Leite, M. Carrington, S. Vaughan, C. Hertz-Fowler, S. Dean, J.D. Sunter, R.J. Wheeler, and K. Gull. 2023. Genome-wide subcellular protein map for the flagellate parasite Trypanosoma brucei. Nat Microbiol. 8:533–547. doi:10.1038/s41564-022-01295-6.

Butenko, A., F.R. Opperdoes, O. Flegontova, A. Horák, V. Hampl, P. Keeling, R.M.R. Gawryluk, D. Tikhonenkov, P. Flegontov, and J. Lukeš. 2020. Evolution of metabolic capabilities and molecular features of diplonemids, kinetoplastids, and euglenids. BMC Biol. 18:23. doi:10.1186/s12915-020-0754-1.

Cavalier-Smith, T. 2016. Higher classification and phylogeny of Euglenozoa. Eur. J. Protistol. 56:250–276. doi:10.1016/j.ejop.2016.09.003.

Chen, G.-Y., C. Deng, D.M. Chenoweth, and M.A. Lampson. 2024. The central spindle drives anaphase chromosome segregation. bioRxiv. 2024.08.30.610502. doi:10.1101/2024.08.30.610502.

Cooke, C.A., M.M. Heck, and W.C. Earnshaw. 1987. The inner centromere protein (INCENP) antigens: movement from inner centromere to midbody during mitosis. J. Cell Biol. 105:2053–2067.

Corkery, D.P., A.C. Holly, S. Lahsaee, and G. Dellaire. 2015. Connecting the speckles: Splicing kinases and their role in tumorigenesis and treatment response. Nucleus. 6:279–288. doi:10.1080/19491034.2015.1062194.

Drinnenberg, I.A., and B. Akiyoshi. 2017. Evolutionary lessons from species with unique kinetochores. Prog. Mol. Subcell. Biol. 56:111–138. doi:10.1007/978-3-319-58592-5_5.

Ebenezer, T.E., M. Zoltner, A. Burrell, A. Nenarokova, A.M.G. Novák Vanclová, B. Prasad, P. Soukal, C. Santana-Molina, E. O’Neill, N.N. Nankissoor, N. Vadakedath, V. Daiker, S. Obado, S. Silva-Pereira, A.P. Jackson, D.P. Devos, J. Lukeš, M. Lebert, S. Vaughan, V. Hampl, M. Carrington, M.L. Ginger, J.B. Dacks, S. Kelly, and M.C. Field. 2019. Transcriptome, proteome and draft genome of Euglena gracilis. BMC Biol. 17:11. doi:10.1186/s12915-019-0626-8.

Faktorová, D., B. Kaur, M. Valach, L. Graf, C. Benz, G. Burger, and J. Lukeš. 2020. Targeted integration by homologous recombination enables in situ tagging and replacement of genes in the marine microeukaryote Diplonema papillatum. Environ Microbiol. 22:3660–3670. doi:10.1111/1462-2920.15130.

Faktorová, D., K. Záhonová, C. Benz, J.B. Dacks, M.C. Field, and J. Lukeš. 2023. Functional differentiation of Sec13 paralogues in the euglenozoan protists. Open Biol. 13:220364. doi:10.1098/rsob.220364.

Flegontova, O., P. Flegontov, P.A.C. Londoño, W. Walczowski, D. Šantić, V.P. Edgcomb, J. Lukeš, and A. Horák. 2020. Environmental determinants of the distribution of planktonic diplonemids and kinetoplastids in the oceans. Environ Microbiol. 22:4014–4031. doi:10.1111/1462-2920.15190.

Herman, J.A., M.P. Miller, and S. Biggins. 2020. chTOG is a conserved mitotic error correction factor. Elife. 9:e61773. doi:10.7554/eLife.61773.

Hirano, T. 2012. Condensins: universal organizers of chromosomes with diverse functions. Genes Dev. 26:1659–1678. doi:10.1101/gad.194746.112.

Hochegger, H., N. Hégarat, and J.B. Pereira-Leal. 2013. Aurora at the pole and equator: overlapping functions of Aurora kinases in the mitotic spindle. Open Biol. 3:120185. doi:10.1098/rsob.120185.

van Hooff, J.J., E. Tromer, L.M. van Wijk, B. Snel, and G.J. Kops. 2017. Evolutionary dynamics of the kinetochore network in eukaryotes as revealed by comparative genomics. EMBO Rep. 18:1559–1571. doi:10.15252/embr.201744102.

Ishii, M., and B. Akiyoshi. 2020. Characterization of unconventional kinetochore kinases KKT10 and KKT19 in Trypanosoma brucei. J Cell Sci. 133:jcs240978. doi:10.1242/jcs.240978.

Kaur, B., M. Valach, P. Peña-Diaz, S. Moreira, P.J. Keeling, G. Burger, J. Lukeš, and D. Faktorová. 2018. Transformation of Diplonema papillatum, the type species of the highly diverse and abundant marine microeukaryotes Diplonemida (Euglenozoa). Environ Microbiol. 20:1030–1040. doi:10.1111/1462-2920.14041.

Kelly, S., J. Reed, S. Kramer, L. Ellis, H. Webb, J. Sunter, J. Salje, N. Marinsek, K. Gull, B. Wickstead, and M. Carrington. 2007. Functional genomics in Trypanosoma brucei: a collection of vectors for the expression of tagged proteins from endogenous and ectopic gene loci. Mol. Biochem. Parasitol. 154:103–109. doi:10.1016/j.molbiopara.2007.03.012.

Komaki, S., E.C. Tromer, G. De Jaeger, N. De Winne, M. Heese, and A. Schnittger. 2022. Molecular convergence by differential domain acquisition is a hallmark of chromosomal passenger complex evolution. Proc Natl Acad Sci U S A. 119:e2200108119. doi:10.1073/pnas.2200108119.

Kostygov, A.Y., A. Karnkowska, J. Votýpka, D. Tashyreva, K. Maciszewski, V. Yurchenko, and J. Lukeš. 2021. Euglenozoa: taxonomy, diversity and ecology, symbioses and viruses. Open Biol. 11:200407. doi:10.1098/rsob.200407.

Laband, K., R. Le Borgne, F. Edwards, M. Stefanutti, J.C. Canman, J.-M. Verbavatz, and J. Dumont. 2017. Chromosome segregation occurs by microtubule pushing in oocytes. Nat Commun. 8:1499. doi:10.1038/s41467-017-01539-8.

Lampson, M.A., and I.M. Cheeseman. 2011. Sensing centromere tension: Aurora B and the regulation of kinetochore function. Trends Cell Biol. 21:133–140. doi:10.1016/j.tcb.2010.10.007.

Lax, G., M. Kolisko, Y. Eglit, W.J. Lee, N. Yubuki, A. Karnkowska, B.S. Leander, G. Burger, P.J. Keeling, and A.G.B. Simpson. 2021. Multigene phylogenetics of euglenids based on single-cell transcriptomics of diverse phagotrophs. Mol Phylogenet Evol. 159:107088. doi:10.1016/j.ympev.2021.107088.

Li, Z., J.H. Lee, F. Chu, A.L. Burlingame, A. Günzl, and C.C. Wang. 2008. Identification of a novel chromosomal passenger complex and its unique localization during cytokinesis in Trypanosoma brucei. PLoS ONE. 3:e2354. doi:10.1371/journal.pone.0002354.

Miller, M.P., C.L. Asbury, and S. Biggins. 2016. A TOG protein confers tension sensitivity to kinetochore-microtubule attachments. Cell. 165:1–12. doi:10.1016/j.cell.2016.04.030.

Musacchio, A., and A. Desai. 2017. A molecular view of kinetochore assembly and function. Biology (Basel). 6:5. doi:10.3390/biology6010005.

Musacchio, A., and E.D. Salmon. 2007. The spindle-assembly checkpoint in space and time. Nat. Rev. Mol. Cell Biol. 8:379–393. doi:10.1038/nrm2163.

Porter, D. 1973. Isonema papillatum sp. n., a new colorless marine flagellate: A light- and electronmicroscopic study. The Journal of Protozoology. 20:351–356. doi:10.1111/j.1550-7408.1973.tb00895.x.

Prokopchuk, G., T. Korytář, V. Juricová, J. Majstorović, A. Horák, K. Šimek, and J. Lukeš. 2022. Trophic flexibility of marine diplonemids - switching from osmotrophy to bacterivory. ISME J. 16:1409–1419. doi:10.1038/s41396-022-01192-0.

Saldivia, M., E. Fang, X. Ma, E. Myburgh, J.B.T. Carnielli, C. Bower-Lepts, E. Brown, R. Ritchie, S.B. Lakshminarayana, Y.-L. Chen, D. Patra, E. Ornelas, H.X.Y. Koh, S.L. Williams, F. Supek, D. Paape, R. McCulloch, M. Kaiser, M.P. Barrett, J. Jiricek, T.T. Diagana, J.C. Mottram, and S.P.S. Rao. 2020. Targeting the trypanosome kinetochore with CLK1 protein kinase inhibitors. Nat Microbiol. 5:1207–1216. doi:10.1038/s41564-020-0745-6.

Schneider, C.A., W.S. Rasband, and K.W. Eliceiri. 2012. NIH Image to ImageJ: 25 years of image analysis. Nat. Methods. 9:671–675.

Schoenle, A., M. Hohlfeld, K. Hermanns, F. Mahé, C. de Vargas, F. Nitsche, and H. Arndt. 2021. High and specific diversity of protists in the deep-sea basins dominated by diplonemids, kinetoplastids, ciliates and foraminiferans. Commun Biol. 4:501. doi:10.1038/s42003-021-02012-5.

Stortz, J.A., T.D. Serafim, S. Alsford, J. Wilkes, F. Fernandez-Cortes, G. Hamilton, E. Briggs, L. Lemgruber, D. Horn, J.C. Mottram, and R. McCulloch. 2017. Genome-wide and protein kinase-focused RNAi screens reveal conserved and novel damage response pathways in Trypanosoma brucei. PLOS Pathogens. 13:e1006477. doi:10.1371/journal.ppat.1006477.

Tashyreva, D., G. Prokopchuk, J. Votýpka, A. Yabuki, A. Horák, and J. Lukeš. 2018. Life Cycle, Ultrastructure, and Phylogeny of New Diplonemids and Their Endosymbiotic Bacteria. mBio. 9:e02447–17. doi:10.1128/mBio.02447-17.

Tashyreva, D., J. Týč, A. Horák, and J. Lukeš. 2023. Ultrastructure and 3D reconstruction of a diplonemid protist (Diplonemea) and its novel membranous organelle. mBio. 14:e0192123. doi:10.1128/mbio.01921-23.

Tashyreva, D., J. Votýpka, A. Yabuki, A. Horák, and J. Lukeš. 2025. Description of new diplonemids (Diplonemea, Euglenozoa) and their endosymbionts: Charting the morphological diversity of these poorly known heterotrophic flagellates. Protist. 177:126090. doi:10.1016/j.protis.2025.126090.

Tedeschi, A., G. Wutz, S. Huet, M. Jaritz, A. Wuensche, E. Schirghuber, I.F. Davidson, W. Tang, D.A. Cisneros, V. Bhaskara, T. Nishiyama, A. Vaziri, A. Wutz, J. Ellenberg, and J.-M. Peters. 2013. Wapl is an essential regulator of chromatin structure and chromosome segregation. Nature. 501:564–568. doi:10.1038/nature12471.

Triemer, R.E. 1992. Ultrastructure of mitosis in Diplonema ambulator Larsen and Patterson (Euglenozoa). Eur J Protistol. 28:398–404. doi:10.1016/S0932-4739(11)80003-9.

Triemer, R.E., and D.W. Ott. 1990. Ultrastructure of Diplonema ambulator larsen & patterson (euglenozoa) and its relationship to Isonema. Eur J Protistol. 25:316–320. doi:10.1016/S0932-4739(11)80123-9.

Tromer, E.C., T.A. Wemyss, P. Ludzia, R.F. Waller, and B. Akiyoshi. 2021. Repurposing of synaptonemal complex proteins for kinetochores in Kinetoplastida. Open Biol. 11:210049. doi:10.1098/rsob.210049.

Valach, M., S. Moreira, C. Petitjean, C. Benz, A. Butenko, O. Flegontova, A. Nenarokova, G. Prokopchuk, T. Batstone, P. Lapébie, L. Lemogo, M. Sarrasin, P. Stretenowich, P. Tripathi, E. Yazaki, T. Nara, B. Henrissat, B.F. Lang, M.W. Gray, T.A. Williams, J. Lukeš, and G. Burger. 2023. Recent expansion of metabolic versatility in Diplonema papillatum, the model species of a highly speciose group of marine eukaryotes. BMC Biol. 21:99. doi:10.1186/s12915-023-01563-9.

de Vargas, C., S. Audic, N. Henry, J. Decelle, F. Mahé, R. Logares, E. Lara, C. Berney, N. Le Bescot, I. Probert, M. Carmichael, J. Poulain, S. Romac, S. Colin, J.-M. Aury, L. Bittner, S. Chaffron, M. Dunthorn, S. Engelen, O. Flegontova, L. Guidi, A. Horák, O. Jaillon, G. Lima-Mendez, J. Lukeš, S. Malviya, R. Morard, M. Mulot, E. Scalco, R. Siano, F. Vincent, A. Zingone, C. Dimier, M. Picheral, S. Searson, S. Kandels-Lewis, Tara Oceans Coordinators, S.G. Acinas, P. Bork, C. Bowler, G. Gorsky, N. Grimsley, P. Hingamp, D. Iudicone, F. Not, H. Ogata, S. Pesant, J. Raes, M.E. Sieracki, S. Speich, L. Stemmann, S. Sunagawa, J. Weissenbach, P. Wincker, and E. Karsenti. 2015. Ocean plankton. Eukaryotic plankton diversity in the sunlit ocean. Science. 348:1261605. doi:10.1126/science.1261605.

Xiang, Y., D. Tsuchiya, Z. Yu, X. Zhao, S. McKinney, J. Unruh, B. Slaughter, C.M. Lake, and R.S. Hawley. 2024. Multiple reorganizations of the lateral elements of the synaptonemal complex facilitate homolog segregation in Bombyx mori oocytes. Curr Biol. 34:352-360.e4. doi:10.1016/j.cub.2023.12.018.

Yatskevich, S., J. Rhodes, and K. Nasmyth. 2019. Organization of Chromosomal DNA by SMC Complexes. Annu Rev Genet. 53:445–482. doi:10.1146/annurev-genet-112618-043633.

Yu, C.-H., S. Redemann, H.-Y. Wu, R. Kiewisz, T.Y. Yoo, W. Conway, R. Farhadifar, T. Müller-Reichert, and D. Needleman. 2019. Central-spindle microtubules are strongly coupled to chromosomes during both anaphase A and anaphase B. Mol Biol Cell. 30:2503–2514. doi:10.1091/mbc.E19-01-0074.

Záhonová, K., J. Lukeš, and J.B. Dacks. 2025. Diplonemid protists possess exotic endomembrane machinery, impacting models of membrane trafficking in modern and ancient eukaryotes. Curr Biol. S0960-9822(25)00195–2. doi:10.1016/j.cub.2025.02.032.

